# Testing for implicit emotion regulation in childhood

**DOI:** 10.1101/2021.12.19.473228

**Authors:** Stepheni Uh, Roma Siugzdaite, Alexander Anwyl-Irvine, Edwin S. Dalmaijer, Giacomo Bignardi, Tess A. Smith, Duncan E. Astle

## Abstract

Although implicit emotion regulation is thought to be critical for psychosocial development and mental wellbeing, few studies have investigated the neural underpinnings of this form of emotion regulation in children. We used a modified emotional Go/NoGo block design fMRI task to explore the neural correlates of implicit emotion regulation and individual differences in a sample of 40 children (50% female, mean age = 8.65 ± 0.77). Conditions included happy, sad, neutral, and scrambled faces as implicit distractors within the actual Go/NoGo targets. We used a relatively standard preprocessing pipeline via fMRIprep, with T-contrasts for response inhibition and emotional effects, and a nonparametric multiple comparisons procedure, with SnPM, for our group-level analysis. There were multiple significant response inhibition effects, including larger NoGo vs Go activation in the IFG, insula, and MCC/ACC. Valence effects showed significantly greater right putamen activity for the Sad NoGo vs Go contrast and greater bilateral putamen and right pallidum activity for the Happy Go vs Sad Go contrast. These results provide preliminary findings of neural substrates, particularly the putamen, that may be associated with implicit emotion regulation in children.

## 1. Introduction

Implicit emotion regulation – that which happens automatically and adaptively without conscious supervision or explicit intentions (Koole & Rothermund, 2011) – has recently garnered theoretical and empirical emphasis. This ability to quickly detect implicit socioemotional cues and automatically regulate both emotional and behavioral responses is particularly critical in children, as they learn to engage in the social world, to develop typical socioemotional functioning, resilience, and mental health (Buckner et al., 2003; Camacho et al., 2019; Urbain et al., 2017). Implicit forms of regulating emotions may also be more representative of realistic interpersonal settings, relative to tasks that require the explicit processing of emotions. For instance, automatically knowing when to engage or inhibit behavioral responses (e.g., converse, withdraw) based on implicit emotional cues (e.g., facial expressions), as well as the ability to self-regulate intrinsic emotional responses to these cues, is constantly required for social interactions and maintaining psychosocial well-being (Todd & Lewis, 2006; Etkin et al., 2011). To date, however, there is a paucity of research investigating the neural bases subserving implicit emotion regulation in children.

### 1.1. Implicit emotion regulation

Much of the emotion regulation literature has characterized the skill as a deliberative and conscious process (Koole & Rothermund, 2011). Recently, researchers have proposed theoretical models separating explicit and implicit types of emotion regulation, albeit with some overlap, based on the nature of the emotion regulation goal as well as the emotion change process (see Braunstein et al., 2017). Implicit emotion regulation may involve bottom-up emotional control networks in situations where emotional information is task-irrelevant (Hung et al., 2018). In this case, these regulatory processes occur at the earliest stages of emotional processing in response to emotional stimuli – potentially without the subjective experience of feeling specific emotions (Ahmed et al., 2015) – and can prevent emotional context from interfering with one’s goals or activities (Urbain et al., 2017). At the neural level, the dorsal anterior cingulate cortex (dACC), dorsolateral and ventrolateral prefrontal regions (dlPFC, vlPFC), as well as the medial prefrontal cortex (mPFC) have been implicated in implicit emotion regulation (Braunstein et al., 2017; Etkin et al., 2015; Gyurak et al., 2011). However, it is still unclear whether these areas are more associated with implicit versus explicit emotion regulation or if they are part of an overarching regulatory network. This lack of clarity is due to overlap of suggested substrates – particularly the lateral prefrontal regions – between both forms of emotion regulation (Gyurak et al., 2011). As studies investigating adaptive and automatic regulatory processes in response to salient stimuli in children are limited (Urbain et al., 2017), the present study focuses specifically on implicit emotion regulation within this age group.

### 1.2. Implicit emotion regulation and socioemotional development

The ability to self-sufficiently regulate emotions is an important socioemotional and cognitive step for children as they grow from their ‘infant phase’ of depending on their caregivers for emotional support (Perlman & Pelphrey, 2011). Implicit emotion regulation strategies may be learned earlier during development, relative to explicit emotion regulation strategies, simply because the latter require more developed cognitive and behavioral regulatory processes (Liberzon et al., 2015). For instance, children learn early on to adapt and control their behaviors and ultimately navigate social interactions when exposed to incidental emotional stimuli, like facial expressions of others (Urbain et al., 2017). In essence, implicit emotion regulation reflects more automatic responses since these processes are activated without explicit instruction (Liberzon et al., 2015). Automatic responding is essential for typical socioemotional development as it provides the capability to rapidly adapt to emotional cues and/or distractors, which can influence socioemotional functioning. Biases for certain implicit emotional cues, moreover, can influence both individual perception of an environment and overall well-being (Surguladze et al., 2005). This is particularly important for children to develop adaptive emotion, as well as self-regulation strategies (Buckner et al., 2003; Hopp et al., 2011; Urbain et al., 2017), which help maintain resilience and psychological health (Hopp et al., 2011). Indeed, the ability to ignore emotional distractors and focus on a task is compromised in most neuropsychiatric and psychological disorders (Bartholomew et al., 2019; Davidson & Slagter, 2000). Furthermore, successful implicit emotion regulation has been linked to reduced depressive symptoms and greater social adjustment (Hopp et al., 2011).

Implicit emotion regulation processes are thus vital for socioemotional development as they operate constantly to prevent emotional context interfering with one’s daily activities (Gyurak et al., 2011; Koole & Rothermund, 2011), to offset the impact of emotional responses triggered automatically from everyday life events (Koole & Rothermund, 2011), and to enhance flexible and efficient action control via psychological adaptation (Gyurak et al., 2011).

### 1.3. Neuroimaging studies of implicit emotion regulation in children

Few neuroimaging studies have examined implicit emotion regulation tasks in childhood. In fact, most task paradigms, like the emotional Go/NoGo task, require explicit emotion regulation, with participants responding directly to an emotional cue such as emotional facial expressions (Urbain et al., 2017). Though, one functional magnetic resonance imaging (fMRI) study assessed implicit emotion regulation with a modified emotional Go/NoGo task in which happy, angry, or neutral faces preceded a non-valence response stimulus to assess inhibition in the face of emotional distractors across age groups of children and young adults (Todd et al., 2012). Children showed greater left orbitofrontal cortex (OFC) activation when inhibiting responses in the Happy condition, compared to the Angry condition. Furthermore, the degree of left OFC activation during the Angry condition increased with age. This pattern aligns with past research indicating that children exhibit a “positivity bias” and first learn and accurately recognize happy facial expressions before sad/angry and then surprise/fear expressions (Camras & Allison, 1985; Herba et al., 2006). Urbain and colleagues (2017), on the other hand, combined this paradigm with magnetoencephalography (MEG) and found that inhibitory responses when exposed to Angry faces recruited the right orbito-frontal gyrus and the left anterior temporal lobe in comparison to Happy faces in children. Other studies that mirrored this emotional Go/NoGo paradigm with aversive stimuli like threat-related images or fearful faces have primarily recruited adolescents in clinical populations (see Brown et al., 2015; Ho et al., 2012) with mixed results. In short, there is a scarcity of evidence of the neural foundations for implicit emotion regulatory processes in children. The present paper provides an opportunity to address this.

### 1.4. Aims

The overarching aim of this study was to investigate the neural substrates of implicit emotion regulation with a modified emotional Go/NoGo fMRI task in a sample of typically developing children. Given the dearth of neuroimaging studies studying implicit emotion regulation in children, we purposely took on an exploratory approach with a whole-brain analysis. This avoids masking off particular areas, making the analysis agnostic as to where effects may emerge. Given the available evidence, we predicted that areas recognized for response inhibition (e.g., dACC, insula, inferior frontal gyrus (IFG) (Hung et al., 2018)) and emotion processing in children (i.e., positivity bias, greater amygdala responses to positive stimuli), as well as the substrates that have been proposed for implicit emotion regulation (e.g., mPFC, dACC, vlPFC) would be key regions reflecting both implicit emotion regulation and response inhibition. A secondary aim was to assess whether these emotional distractors would still elicit valence-specific effects. Thirdly and finally, as implicit emotion regulation has been linked to greater psychological health and reduced depressive symptoms (Hopp et al., 2011), we tested for potential relationships between neural responses of implicit emotion regulation and individual differences in mental wellbeing and emotional and behavioral difficulties.

## 2. Methods

### 2.1. Participants and procedure

Participants for this study (n = 70) were recruited via posters, word of mouth, and online Facebook advertisements throughout Cambridge, United Kingdom. All participants underwent a medical screening for MRI scanning eligibility prior to participating in the study and provided informed, written consent. Ethical approval was granted by the University of Cambridge Psychology Research Ethics Committee (PRE.2017.102). Thirty participants were excluded for excessive movement (average framewise displacement > 0.5 mm), signal loss, and insufficient scan time. The final sample, therefore, included 40 participants (50% female) aged 7-9 years old (M = 8.65; SD = 0.77) who completed the full emotional Go/NoGo fMRI task.

### 2.2. Emotional go/nogo task

The design of this Emotional Go/NoGo task was modelled on a paradigm used by Ho and colleagues (2012), focusing on the implicit emotional distractors. These distractors were presented on every trial and included happy, sad, or neutral facial expressions and a ‘cold cognition’ (control) condition that matched the basic stimulus properties (size, shape, luminosity) without a discernable face. Children were told to attend to the shape (circle or square) that appeared overlaid on the face. In the current study, the circle and square served as the Go/NoGo stimuli and were counterbalanced across participants, to control for any perceptual differences between the two. Participants were instructed to press a button with the index finger of their dominant hand to the Go trials only. As our main interest is implicit emotion regulation, we adapted Ho and colleagues’ (2012) block design with Go blocks consisting of only Go trials and NoGo blocks consisting of 12 Go and 6 NoGo trials (both circles and squares). Blocks were separated by emotional condition (e.g., Happy Go block, Happy NoGo block, etc.) for robust data acquisition. We purposely used a block design to maximize detection power (Shan et al., 2014) in consideration of our sample’s age range and propensity for head motion in the scanner (Yuan et al., 2008).

The facial expressions for the emotional conditions were selected from the Dartmouth Database of Children’s Faces (Dalrymple et al., 2013). Due to evidence indicating the strong influence of peer relations/friendships on adolescent mental health, cognition, and behavior (Blakemore & Mills, 2014), grey-scaled child faces were used for the task conditions. In total, 18 frontal facial identities were selected (nine girls, nine boys) – each expressing happy, sad, and neutral – based upon perceived age to match our sample of 7–9-year-old children as well as highest rankings for emotion validation scored by independent raters from Dalrymple and colleagues’ (2013) study. A scrambling algorithm was used to produce scrambled faces without any discernible facial features or emotional valence, while maintaining the luminance distribution, for the control condition. Only pixels containing the child’s face were scrambled, which was detected using a luminance threshold, thereby ensuring that a black background remained intact. Each ‘block’ was 6×6 pixels and was shuffled without replacement.

There were two task runs, each consisting of 8 blocks: 4 Go blocks and 4 NoGo blocks, separated by emotional condition. Each block has 18 trials for a total of 288 trials across the two runs. Within a NoGo block, there are 12 Go trials (e.g., circles) and 6 No-Go trials (e.g., squares) to induce the prepotent tendency to respond. The facial identity for each trial is shown 250-550 ms prior to the presentation of the shapes. Once the shape appears, a response time window is set for a maximum of 1000 ms followed by a 1000 ms fixation cross. Overall, the inter-trial interval (ITI) is jittered between 1250-1750 ms, and the maximum duration of the entire task is 20 minutes. When a participant responds, the shape turns grey to indicate that their response has been recorded. There is also an optional 1000 ms break between blocks during which the instructions are displayed again, and participants are instructed to press the button if they are ready to continue. The order in which the block conditions are presented in addition to the order of the trials within each block are randomized.

The stimuli were presented and data were recorded using PyGaze (Dalmaijer et al., 2014) and PsychoPy 2 (Peirce et al., 2019). The scripts for generating scrambled faces, randomization and stimulus presentation are available here: https://github.com/u01ai11/emoFace/

### 2.3. Individual differences

To explore individual differences across mental health as well as emotional and behavioral challenges in relation to neural responses from the emotional Go/NoGo task, we administered the RCADS short form subscales for anxiety (GAD) and depression (MDD) (Muris et al., 2002) and the Strengths and Difficulties Questionnaire (SDQ) (Goodman, 1997). The SDQ was completed by caregivers while the RCADS subscales were self-reported by the participants.

### 2.4. fMRI data acquisition and preprocessing

#### Image Acquisition

MRI scans were obtained on a 3T Siemens Prisma with a 32-channel quadrature head coil. Before the scan, all children practiced going into a realistic mock scanner and lying still to minimize movements. For the Emotional Go/NoGo task, T2*-weighted fMRI data was acquired using a Gradient-Echo Echo-Planar Imaging (EPI) sequence. The number of volumes acquired varied depending on participants’ response times to trials and acquisition time ranged from approximately 15-20 minutes with the following parameters: Repetition Time (TR) = 2000 ms; Echo Time (TE) = 30 ms, flip angle = 78 degrees; voxel dimensions = 3 mm isotropic; FOV = 192 × 192 mm; acquisition matrix = 64 × 64; number of slices = 32; descending sequential sequence. For registration of the functional images, a high-resolution T1-weighted (T1w) structural image was acquired over 4 minutes 32 seconds using a Magnetization Prepared Rapid Gradient Echo (MPRAGE) sequence (TR = 2250 ms; TE = 3.02 ms; Inversion Time (TI) = 900 ms; flip angle = 9 degrees; number of slices = 192; voxel dimensions = 1 mm^3^ isotropic; GRAPPA acceleration factor = 2).

#### Image preprocessing

Preprocessing was performed using *fMRIPrep* 1.5.0 (Esteban, Markiewicz, et al., 2019) based on *Nipype* 1.2.2 (Gorgolewski et al., 2011). All raw DICOM files were first converted into the Brain Imaging Data Structure (BIDS) format and then preprocessed using the fMRIPrep workflow (see Esteban et al., 2019). This included skull-stripping and anatomical reconstruction to standard space of the T1w image and the following fMRI data preprocessing steps: realignment (head-motion estimation and correction using MCFLIRT, FSL); slice-timing correction (AFNI); boundary-bound co-registration of the fMRI data to the T1w reference image (Greve and Fischl, 2009), configured with six degrees of freedom; resampling and spatial normalization into MNI152 standard space (ICBM 152 Nonlinear Asymmetrical template version 2009c; Fonov et al., 2009). Several confounding time-series were calculated based on the preprocessed images: framewise displacement (FD), DVARS and three region-wise global signals. FD and DVARS are calculated for each functional run, both using their implementations in *Nipype* (following the definitions by Power et al., 2014). Children were excluded for high average motion (mean FD > 0.5 mm or presence of major motion artifacts, n = 30).

### 2.5. Statistical analysis

#### Task behavioral measures

D-prime, a sensitivity measure capturing response bias, was calculated to account for hits (correct response to Go trials) and false alarms (commission errors or incorrect response to NoGo trials) across each emotional condition (happy, sad, neutral, control) (Stainslaw & Todorov, 1999; Wiemers & Redick, 2019). D-prime is computed by subtracting the z-scored hit rate by the z-scored false alarm rate. Hit rates for each emotional condition were calculated by subtracting the number of hits by the number of omission errors (no response to Go trial) then dividing this value by the number of total Go trials. A logarithmic adjustment (adding 0.5 to hits – omission errors and adding 1 to the number of Go trials) was made to ensure that the d-prime calculation would work even if participants did not make any errors (see Wiemers & Redick 2019). The false alarm rate was calculated with the same logarithmic adjustment. Average response times for Go trials only within Go blocks as well as within NoGo blocks across each emotional condition were also computed.

#### fMRI analysis

Statistical Parametric Mapping (SPM12; University College, London, UK) software with *Nipype* was used to implement the first level analysis workflow (see Esteban et al., 2020) to identify blood-oxygen-level-dependent (BOLD) responses related to emotional (e.g., Happy, Sad, Neutral blocks) and cognitive effects (Go and NoGo blocks). Prior to model estimation, the functional images were spatially smoothed using a Gaussian kernel (6 mm FWHM) and filtered with a high pass filter (128 s). As a blocked design, BOLD responses were modeled at each block onset and then convolved with a standard hemodynamic response function (HRF) and its temporal derivative producing a general linear model (GLM) for each participant. The GLM included six confounding regressors representing the movement parameters derived from the realignment stage to correct for head movement. We conducted T-contrasts looking at the main effects of response inhibition (Go vs NoGo), facial effect (Faces vs Cold Cognition), emotional response inhibition (Sad Go vs Sad NoGo, Happy Go vs Happy NoGo), valence effects (Sad vs Happy, Sad vs Neutral, Happy vs Neutral), and also valence effects in Go blocks only (Sad Go vs Happy Go, Sad Go vs Neutral Go, Happy Go vs Neutral Go) as no inhibition would be required and button pressing would likely be a passive response.

In order to combat the multiple comparisons issue, we utilized a nonparametric multiple comparisons procedure based on permutation testing for our group-level analysis using SnPM13.1.06 software with SPM12 in MATLAB 2018b (see htpp://nisox.org/Software/SnPM13/). Through this toolbox, we were able to run multi-subject one sample T-tests on each contrast by using the GLM to construct pseudo *t*-statistic images and then assess those for significance against the null hypothesis generated from this nonparametric resampling-based approach (Nichols & Holmes, 2001). The null distribution is formed by randomly flipping the sign of each participant’s contrast across a number of iterations or permutations. For cluster-wise inference, we ran 5000 cluster-based permutations with a smoothed variance of 6 mm FWHM to match the within subject smoothing and with *p* < 0.05 FWE corrected, using a cluster-forming threshold (CFT) of *p* = 0.0001. CFTs can be statistically arbitrary, and they often reflect unique aspects of the data (Friston et al., 1994; Woo et al., 2014). Having a more stringent threshold (i.e., *p* < 0.001 or *p* < 0.0001), however, is particularly important for studies with smaller sample sizes as this reduces the potential of false positives and increases sensitivity for making inferences about the locations of activation clusters (Woo et al., 2014). Nonparametric permutation methods for neuroimaging data, furthermore, provide an advantageous method to correct for multiple comparisons while requiring minimal assumptions (Eklund et al., 2016; Nichols & Holmes, 2001).

To perform correlations with behavioral measures, masks were then created by extracting the significant clusters from our thresholded activation map of each contrast. Regions of significant clusters were identified from these thresholded images for areas that were not labeled as NaN. Each participant’s average beta values within these significant clusters were then computed from their first level contrasts to run correlations with our measures of interest for individual differences.

#### Individual differences

Kendall’s rank pairwise correlations were run between average beta values from significant areas of activation of specific contrasts from the final 40 participants with measures reflecting mental health (RCADS MDD, RCADS GAD) and emotional and behavioral problems (SDQ total score, SDQ internalizing score, SDQ externalizing score). As the pairwise correlations were run between complete observations, the total number of pairwise comparisons varied across the sample as some participants had missing data (see Results). Kendall’s rank correlations were run for more robust analysis as we have a sample size n < 50 (Bonett & Wright, 2000).

## 3. Results

### 3.1. Task behavioral results

D-prime mean (M) and standard deviation (SD) values were as follows: Happy, M = 2.86, SD = 0.86; Sad, M = 2.67, SD = 0.97; Neutral, M = 2.65, SD = 0.94; Control, M = 2.67, SD = 0.92. There was no significant difference in average d-prime values across all conditions (*F*_3, 117_ = 1.145, *p* = 0.334).

A Kenward-Roger corrected type II ANOVA with fixed factors of emotional condition type (happy, sad, neutral, control) and block type (Go, NoGo) and within subjects as the random factor did not show a significant interaction between condition and block types (*F*_3, 9250_ = 1.45, *p* = 0.23) on average response time for Go trials. There was no main effect of emotion condition (*F*_3, 9250_ = 2.11, *p* = .10), but block type had a significant main effect on average response time for Go trials (*F*_1, 9250_ = 184.77, *p* < .0001). This is because, unsurprisingly, response times to go trials within Go blocks were significantly faster than response times to Go trials within NoGo blocks across each emotional condition (all *p*s < 0.0001, Bonferroni-holm adjusted for multiple comparisons) (Figure 1).

**Figure 1.**
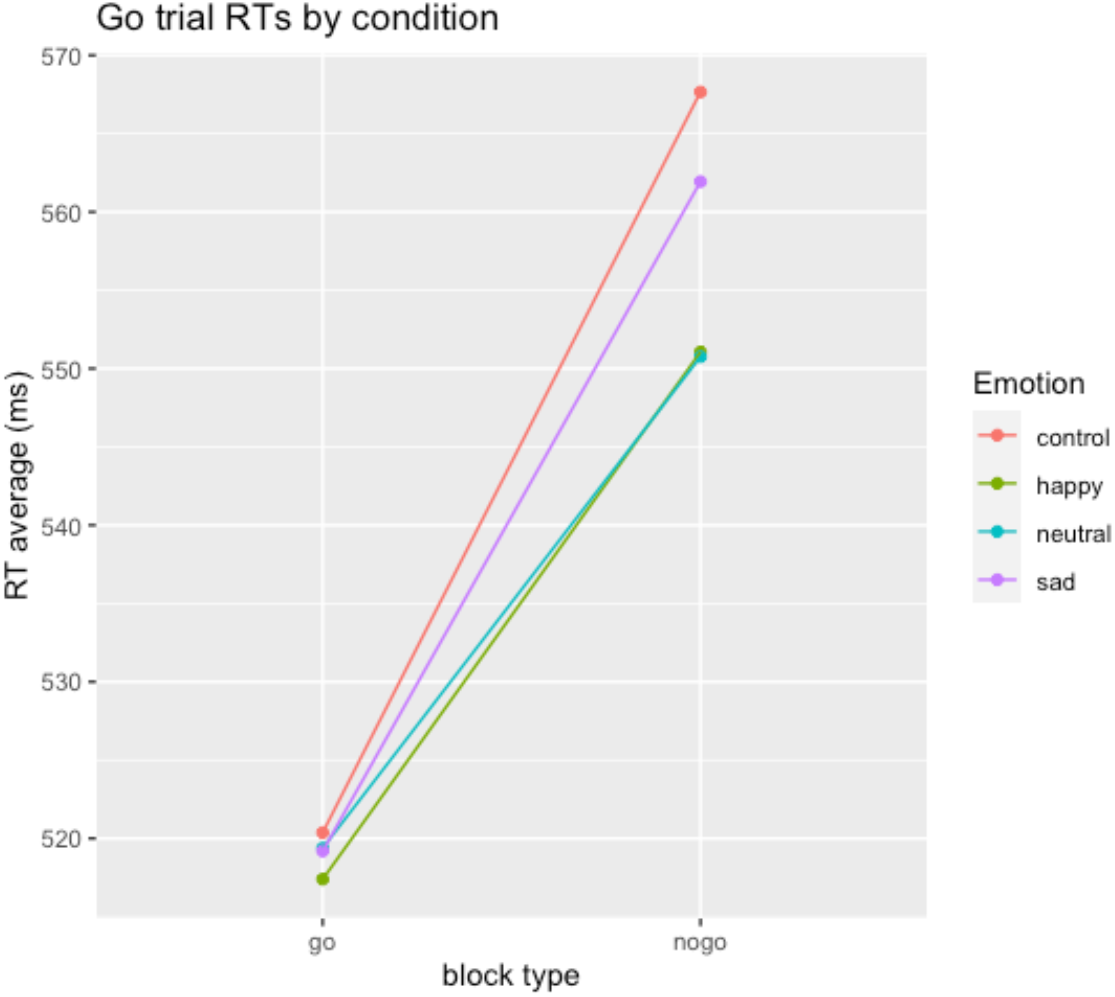
Average response times (RT) in ms to go trials between block types across the four conditions. Within each emotional condition, average response times to go trials in the Go blocks were significantly faster than average response times to go trials in NoGo blocks

### 3.2. fMRI results

fMRI results are listed in Table 1. Main effects were assessed to validate whether the emotional Go/NoGo task elicits the expected neural activations that have been strongly associated with response inhibition and facial stimuli. In the response inhibition contrasts, we found larger NoGo vs Go activation in the vlPFC, specifically in the inferior frontal gyrus (IFG), part of the insula, and the middle and anterior cingulate cortex (MCC/ACC) (Figure 2A). In the Faces vs Control contrast, we found greater activation in the inferior orbital gyrus (IOG)/fusiform gyrus (FFG) (Figure 2B). The Sad NoGo vs Sad Go contrast, which was the only emotional response inhibition contrast that elicited neural activity, showed a significant difference in the right putamen (Figure 2C). Finally, when looking at only Go blocks for emotion effects, the Happy Go vs Sad Go contrast revealed greater activation in the right putamen and pallidum (Figure 2D). Contrasts subtracting emotional as well as neutral blocks from the control condition (scrambled faces) elicited similar activations in the bilateral occipital lobes across all contrasts except for the happy condition (see Supplementary Table 1).

**Table 1.** Main effects, emotional response inhibition (RI), and emotion fMRI contrasts results indicating side of hemisphere (R = Right; L = Left), *p*-value, FWE-corrected at *p* < 0.05, cluster size (*k*), and MNI coordinates XYZ (mm) for cluster peak. IFG = inferior frontal gyrus. MCC = middle cingulate cortex. ACC = anterior cingulate cortex. IOG = inferior orbital gyrus. FFG = fusiform gyrus.

| fMRI Contrast Results |  |  |  |  |  |  |  |  |
| --- | --- | --- | --- | --- | --- | --- | --- | --- |
|  |  | Region | Side | pFWEcorr | k | X | Y | Z |
| Contrast |  |  |  |  |  | (mm) |  |  |
|  | Main |  |  |  |  |  |  |  |
|  | NoGo > Go | IFG, triangular part/ | R | 0.0038 | 66 | 48 | 18 | -3 |
|  |  | Insula/ | R |  |  |  |  |  |
|  |  | IFG, opercular part/ | R |  |  |  |  |  |
|  |  | IFG pars orbitalis | R |  |  |  |  |  |
|  |  | MCC/ | R | 0.0448 | 14 | 6 | 27 | 31 |
|  |  | ACC | R |  |  |  |  |  |
|  | Faces > Control | IOG/ | L | 0.0434 | 16 | -39 | -81 | -7 |
|  |  | FFG | L |  |  |  |  |  |
|  | Valence, Go/NoGo |  |  |  |  |  |  |  |
|  | Sad NoGo > Sad Go | Putamen | R | 0.0162 | 16 | 21 | 6 | 8 |
|  | Valence |  |  |  |  |  |  |  |
|  | Happy Go > Sad Go | Putamen | L | 0.0022 | 102 | -24 | 12 | -7 |
|  |  | Putamen/ | R | 0.0042 | 74 | 24 | 6 | 5 |
|  |  | Pallidum | R |  |  |  |  |  |

**Figure 2.**
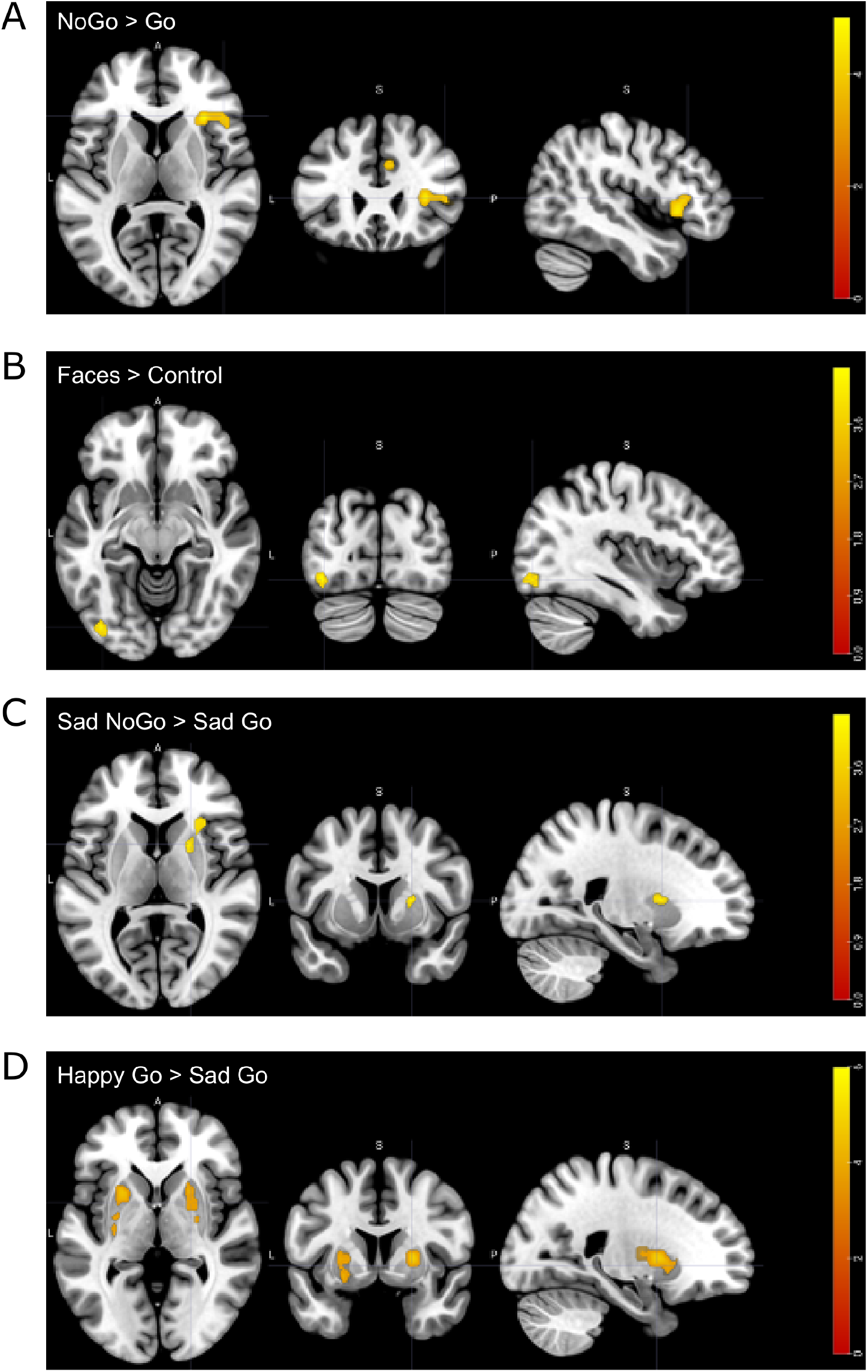
Significant fMRI main effects, emotional response inhibition, and emotion contrast results: A) NoGo > Go showing greater activation in right IFG/insula and right MCC/ACC; B) Faces > Control showing greater activation in IOG/FFG for conditions with facial stimuli; C) Sad NoGo > Sad Go eliciting greater activation in right putamen; D) Happy Go > Sad Go showing greater activation in bilateral putamen and right pallidum

### 3.3. Individual differences results

Significant relationships between all individual differences measures and beta values from the significant activations of the main effects (NoGo > Go, Faces > Control), emotional response inhibition (Sad NoGo > Sad Go), and emotion (Happy Go > Sad Go) contrasts with our measures of individual differences are shown in Figure 3. The strongest relationship identified was a negative correlation between the Sad NoGo > Sad Go beta values and self-reported anxiety (RCADS GAD), τ_b_ = -0.22, p = 0.05. However, this relationship did not survive after correcting for multiple comparisons through the false discovery rate (FDR) correction approach.

**Figure 3.**
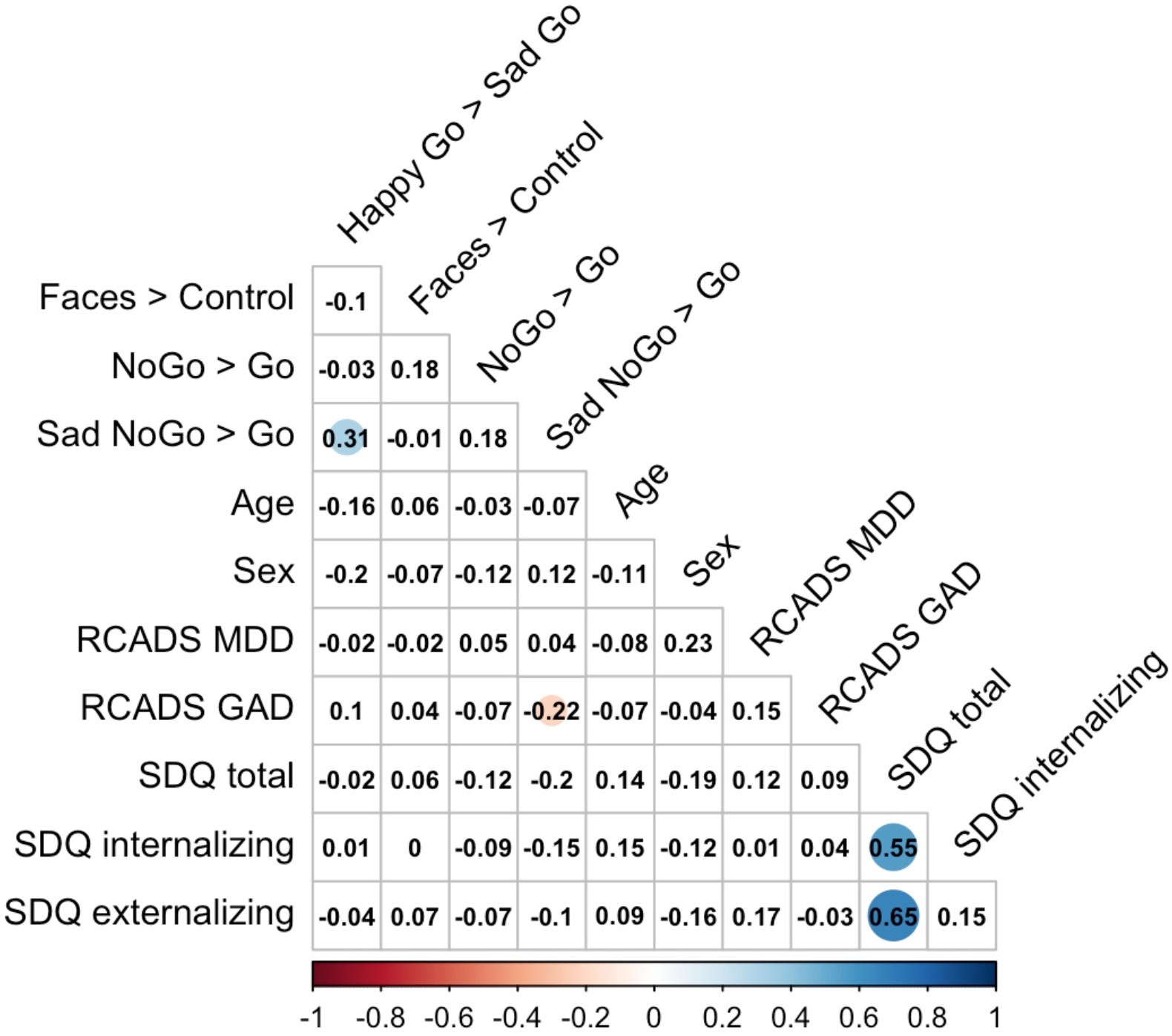
Correlation matrix of all variable intercorrelations highlighting τ_b_ coefficient values of significant correlations (*p*s < 0.05) before multiple comparison corrections

Due to several participants missing some of their data, the number of pairwise comparisons varied from 36-40 (Table 2). The number of complete observations for each variable as well as their means and standard deviations are listed in Supplementary Table 2.

**Table 2.** Number of participants for pairwise comparisons excluding missing data across all variables.

| Variable | 1 | 2 | 3 | 4 | 5 | 6 | 7 | 8 | 9 | 10 | 11 |
| --- | --- | --- | --- | --- | --- | --- | --- | --- | --- | --- | --- |
| 1 Happy Go > Sad Go | 40 |  |  |  |  |  |  |  |  |  |  |
| 2 Faces > Control | 40 | 40 |  |  |  |  |  |  |  |  |  |
| 3 NoGo > Go | 40 | 40 | 40 |  |  |  |  |  |  |  |  |
| 4 Sad NoGo > Go | 40 | 40 | 40 | 40 |  |  |  |  |  |  |  |
| 5 Age | 38 | 38 | 38 | 38 | 38 |  |  |  |  |  |  |
| 6 Sex | 40 | 40 | 40 | 40 | 38 | 40 |  |  |  |  |  |
| 7 RCADS MDD | 38 | 38 | 38 | 38 | 36 | 38 | 38 |  |  |  |  |
| 8 RCADS GAD | 38 | 38 | 38 | 38 | 36 | 38 | 38 | 38 |  |  |  |
| 9 SDQ total | 38 | 38 | 38 | 38 | 38 | 38 | 36 | 36 | 38 |  |  |
| 10 SDQ internalizing | 38 | 38 | 38 | 38 | 38 | 38 | 36 | 36 | 38 | 38 |  |
| 11 SDQ externalizing | 38 | 38 | 38 | 38 | 38 | 38 | 36 | 36 | 38 | 38 | 38 |

## 4. Discussion

The primary aim of this study was to test for implicit emotion regulation within the neural responses of typically developing children and establish whether valence effects are elicited by the modified emotional Go/NoGo task. This modified Go/NoGo task reflects processes that may be required for everyday social interactions and exposures to salient stimuli. For instance, when interacting with others, we are often exposed to micro-expressions or emotional responses that we learn to quickly detect and adapt, often without conscious awareness, our own behaviors to match the social setting (Urbain et al., 2017). However, as implicit emotion regulatory processes at the neural level are relatively unknown, particularly at a young age, our primary focus was to first establish the neural substrates underlying this specific domain of emotion regulation. We then assessed whether this task design elicits valence responses and also explored potential effects of individual differences including mental wellbeing and behavioral challenges.

### 4.1. Behavioral performance

Introducing emotional stimuli as distractors rather than the actual target of the task provides a different representation of implicit emotion regulation. It also allows for the processes by which emotions relate to cognition and behavior – and ultimately developmental outcomes – to be explored (Cole et al., 2004; Ho et al., 2012). Overall, the inhibitory effect was only seen on the average response times since children were significantly faster in responding to Go trials in Go blocks in comparison to Go trials in NoGo blocks across all conditions. There were no significant valence effects on behavioral responses. For example, children did not have more difficulty inhibiting their responses to the happy compared to the sad implicit distractors. However, one possibility for this result is that our study may be limited in power due to our final sample size.

### 4.2. Neural substrates: main task effects

Due to the limited number of studies investigating neural underpinnings of implicit emotion regulation in children, we first assessed whether this modified emotional Go/NoGo task would elicit well-established areas associated with response inhibition. The response inhibition contrast of NoGo > Go elicited activity in areas previously associated with inhibitory processes including the right IFG, insula, and ACC. The IFG has been consistently linked to behavioral impulse control such as response inhibition (Aron et al., 2004; Mitchell, 2011) as well as modulating emotional responses (Mitchell, 2011). Some have even suggested the right IFG is specialized for inhibition alone (Aron et al., 2004); though others have shown that the right IFG is also recruited during phases of attentional control and when detecting important cues (Hampshire et al, 2010). The insula has also been implicated in inhibitory control, with the right insula specifically associated with detecting behaviorally salient events (Cai et al., 2014). The insula is also known to be functionally connected with the ACC (Cai et al., 2014), an area important for conflict monitoring and error detection (Brown & Braver, 2005). As prefrontal and subcortical maturation takes time to develop (Casey et al., 2019), it is possible that these areas are primarily important for early inhibitory processes in children. Our finding of the NoGo blocks eliciting greater activity in the right IFG, insula, and MCC/ACC suggest that our modified blocked emotional Go/NoGo paradigm works to reveal response inhibition circuitry in childhood.

### 4.3. Neural substrates: implicit emotional effects

Although there were no apparent emotion effects at the behavioral level, we investigated whether there would be differential valence responses at the neural level. To explore emotion effects of the implicit emotional distractors, we looked at NoGo vs Go contrasts for each emotional condition as well as overall emotional contrasts of the Go blocks only, since these blocks did not require inhibition. The only condition that displayed a difference in brain activity between the NoGo and Go contrasts was the sad condition. The Sad NoGo blocks compared to the Sad Go blocks elicited greater activation in the right putamen. The role of the putamen in response to emotional cues is relatively unclear (Stuhrmann et al., 2011), especially in younger cohorts, but it has been associated with responses to negative facial stimuli. In clinical samples, for instance, the putamen has been implicated in having a role in hyperactivation to negative stimuli in major depression (Stuhrmann et al., 2011).

As the NoGo blocks involve the engagement of inhibitory behavior, we also exclusively looked at Go blocks to investigate whether these blocks would display more valence-related brain activity. Responding in these blocks could potentially reflect more passive processing, requiring less cognitive load, which may then allow for the emotional distractors to have a stronger impact. Consistent with prior studies looking at age effects of negative emotional cues, the Happy Go blocks elicited bilateral putamen as well as right pallidum activity, in comparison to Sad Go blocks. Specifically, these results align with Todd et al.’s (2011) study in which positive stimuli elicited greater putamen activation than negative stimuli in children while the activity was reversed for adults. Our finding suggests that greater putamen activity in response to positively-valenced cues could be age-specific, as subcortical regions continue to develop and mature with cortical areas in children (Casey et al., 2019). Furthermore, the ventral pallidum has been implicated as a key limbic connection for reward processing (Smith et al., 2009), so our results may reflect a positivity bias associated with children.

### 4.4. Individual difference

The ability to regulate emotions and behaviors is linked to a variety of internal and external influences and is particularly at risk in those who face mental health (Gross & McRae, 2020). Even at a young age, emotion dysregulation has been associated with adolescent depression and anxiety (Young et al., 2019) and is also a known risk factor for developmental and other mental health challenges like self-harm (Uh et al., 2021). Thus, we explored whether our neuroimaging findings would vary based upon individual differences reflecting internal and external problems as well as anxiety and depressive symptoms. No significant relationships were found upon correcting for multiple comparisons, suggesting that the activity in the neural substrates did not vary across these factors of individual differences. However, this could be due to the limitation we faced with power in consideration of our final sample size after preprocessing the imaging data.

### 4.5. Limitations

The present study has several limitations worth noting. First, the sample size was relatively small, which was due to the artifacts present in many of the participants’ fMRI scans. This means that there could be more subtle, but meaningful, valence effects that we are simply unable to detect. Second, the block design was used to maximize robustness and detection power at the potential cost of estimation efficiency or individual responses to specific trials (Shan et al., 2014). However, we utilized a nonparametric permutation approach to analyze whole-brain activity, which minimized assumptions while approaching the multiple comparisons issue present in neuroimaging analyses. Third, we did not have a specific test of how implicit this task really is and thus there is the potential of blurring the boundary between explicit and implicit emotion regulation. On the other hand, there is limited knowledge as to what underlies implicit emotion regulation at the neural level in addition to the likely ‘porous’ boundaries between implicit and explicit domains of emotion regulation, making it difficult to actually compare and test the true implicitness of these tasks (Braunstein et al., 2017). Lastly, we did not collect data from the participants on whether they recognized the emotional expressions used in the study as the implicit emotion distractors. Though, we attempted to capture the highest salience by only selecting expressions that have been rated in the top 90^th^ percentile of accurate emotion recognition by raters in Dalrymple and colleagues’ (2013) study.

### 4.6. Conclusion

To date, few studies have investigated implicit emotion regulation in children, which is thought to be an important capability for overall wellbeing and development. Our preliminary findings from a modified implicit emotion regulation task provide a window into possible neural underpinnings of implicit emotion regulation in a sample of typically developing children. Although we found no behavioral differences in the Go/NoGo task itself, the right putamen showed greater activity for the Sad NoGo blocks in comparison to the Sad Go blocks. The putamen and right pallidum, moreover, may subserve the positivity bias in children as these areas were activated more strongly for the Happy Go condition relative to the Sad Go condition. The task design of our study, furthermore, seems to engage well-established response inhibitory circuitry in a children sample. Future studies investigating this task with a larger sample size and different emotional conditions would be important to further gauge the key substrates for implicit emotion regulation – as well as potential relationships with individual differences in mental health – to better understand what may underlie implicit emotion regulation and its role in psychosocial development.

