## Supplemental Table 1 and Supplemental Table 2 for "Testing for implicit emotion regulation in childhood"

### Testing implicit emotion regulation in childhood: Supplementary material

Emotion fMRI contrasts with the control condition

| | | | Region | Side | pFWEcorr | k | X | Y | Z |
|---|---|---|---|---|---|---|---|---|---|
| **Contrast** | | | | | | | | (mm) | |
| | **Valence** | | | | | | | | |
| | | Happy Go > Control Go | Putamen | L | 0.002 | 80 | -30 | -15 | -4.5 |
| | | | IOG/MOG | L | 0.0112 | 37 | -42 | -7 | -3 |
| | | | MTG | L | 0.0114 | 36 | -48 | -45 | 5 |
| | | | IOG | R | 0.0248 | 23 | 39 | -87 | -6.75 |
| | | | ACC | L | 0.0248 | 23 | -12 | 27 | 23 |
| | | | MFG/SFGmedial | L | 0.0244 | 19 | -27 | 42 | 23 |
| | | Control Go > Neutral Go | Occipital pole | R | 0.012 | 32 | 12 | -102 | -3 |
| | | | Occipital pole | L | 0.019 | 23 | -15 | -102 | -6.75 |
| | | Control Go > Sad Go | Occipital pole | R | 0.0106 | 34 | 12 | -102 | -3 |

***Supplementary Table 1.*** Emotion fMRI contrasts with the control condition (scrambled faces) that elicited significant neural activity results, indicating side of hemisphere (R = Right; L = Left), $p$-value, FWE-corrected at $p < 0.05$, cluster size ($k$), and MNI coordinates XYZ (mm) for cluster peak. IOG/MOG = inferior/middle orbital gyrus. MTG = middle temporal gyrus. ACC = anterior cingulate cortex. MFG/SFGmedial = middle/superior frontal gyrus. *Note: Control Go > Happy Go did not elicit any significant neural activity

| | Variable | N | Mean | SD |
|---|---|---|---|---|
| Beta values | | | | |
| | NoGo > Go | 40 | 0.462 | 0.259 |
| | Sad NoGo > Sad Go | 40 | 0.647 | 0.306 |
| | Faces > Control | 40 | 2.23 | 1.286 |
| | Happy Go > Sad Go | 40 | 0.474 | 0.281 |
| Mental health | | | | |
| | RCADS GAD | 38 | -0.353 | 0.985 |
| | RCADS MDD | 38 | -0.316 | 0.897 |
| Emotional/behavioral problems | | | | |
| | SDQ externalizing | 38 | 5.579 | 4.291 |
| | SDQ internalizing | 38 | 4.447 | 3.644 |
| | SDQ total | 38 | 9.868 | 6.321 |

***Supplementary Table 2.*** Number of complete observations, means, and standard deviations (SD) for each variable
